# Defining the relevant combinatorial space of the PKC/CARD-CC signal transduction nodes

**DOI:** 10.1101/228767

**Authors:** Jens Staal, Yasmine Driege, Mira Haegman, Marja Kreike, Styliani Iliaki, Domien Vanneste, Inna Affonina, Harald Braun, Rudi Beyaert

## Abstract

Biological signal transduction typically display a so-called bow-tie or hour glass topology: Multiple receptors lead to multiple cellular responses but the signals all pass through a narrow waist of central signaling nodes. One such critical signaling node for several inflammatory and oncogenic signaling pathways in humans are the CARD-CC / Bcl10 / MALT1 (CBM) complexes, which get activated by upstream protein kinase C (PKC). In humans, there are four phylogenetically distinct CARD-CC family (CARD9, −10, −11 and −14) proteins and 9 true PKC isozymes (α to ι). At this moment, less than a handful of PKC/CARD-CC relationships are known from experimental evidence. In order to explore the biologically relevant combinatorial space out of all 36 potential permutations in this two-component signaling event, we made use of CRISPR/Cas9 genome-edited HEK293T cells to mutate CARD10 for subsequent pairwise cotransfections of all CARD-CC family members and activated mutants of all true PKCs. By quantitative reporter gene expression readout, we could define specific strong and weak PKC/CARD-CC relationships. Surprisingly as many as 21 PKC/CARD-CC combinations were found to have synergistic effects. We also discovered heterodimerization between different CARD-CC proteins, and that this can influence their PKC response profile. This information will be valuable for future studies of novel signaling pathways dependent on the CBM complex signaling nodes.

## Introduction

Signal transduction aims to transmit a signal from an external or internal stimulus resulting in a cellular response. One such example are the multiple innate and adaptive immune pathways leading to NF-κB activation [1]. Since inappropriate activation can be detrimental, a general problem in cellular signal transduction is that signals and responses to stimuli need to be tightly regulated in order to be sensitive enough while still filtering out environmental noise that could lead to inappropriate activation [2–5].

A common network topology for intracellular molecular signaling pathways is the so-called bow-tie (or hourglass) architecture: multiple receptors and environmental signals pass their information through a few common signaling nodes but end up with a wide diversity of cellular (*i.e*. transcriptional) responses [6,7]. This narrow waist line network topology is thought to contribute to the filtering of informative signals from noise and to ensure appropriate cellular responses [8–10].

One class of such narrow-waist signaling nodes is represented by the CARD-CC/Bcl10/MALT1 (CBM) signaling complexes. Several different CBM signaling complexes exist, which are composed of Bcl10, MALT1 and specific CARD-CC family [11] proteins (CARD9 [12], CARD11 (also known as CARMA1) [13], CARD14 (also known as CARMA2) [14–16] and CARD10 (also known as CARMA3) [17]; Fig. 1) and which are formed upon stimulation of distinct receptors in several immune and non-immune cells. Part of the CBM complex specificity comes from cell type specific expression of the different CARD-CC proteins: in general, CARD9 is expressed in myeloid cells, CARD11 in lymphocytes, CARD14 in skin and CARD10 in most of the other cells [18,19]. Due to their cell-type specific expression, the different CARD-CC proteins play different roles. For example: Loss of *CARD9* function causes susceptibility to fungal infections due to defective innate immune responses [12]. Loss of *CARD11*, on the other hand, cause severe deficiencies in adaptive immunity [20], whereas hyper-active mutants cause B cell lymphomas or B cell Expansion with NF-κB and T cell Anergy (BENTA) [21–23]. The role of *CARD14* is currently unknown, but activating mutations in this CARD-CC have been shown to cause psoriasis or *pityriasis rubra pilaris* (PRP) [24], and elevated *CARD14* expression has been found in an androgen-insensitive and highly tumorogenic prostate cancer sub-clone [25]. Inactivating mutations in *CARD14* are associated to the development of atopic dermatitis [26,27], indicating that CARD14 plays an important role of the skin innate immunity and barrier functions. There are no known human loss- or gain-of-function mutants in *CARD10*, but there are genetic associations to primary open angle glaucoma (POAG) and mild cognitive impairment [28,29]. The *Card10* mouse knock out mutant is often embryo lethal since half of the embryos die from neural tube defects [30]. On the other hand, some cancers show elevated *CARD10* expression [31,32].

**Fig. 1:**
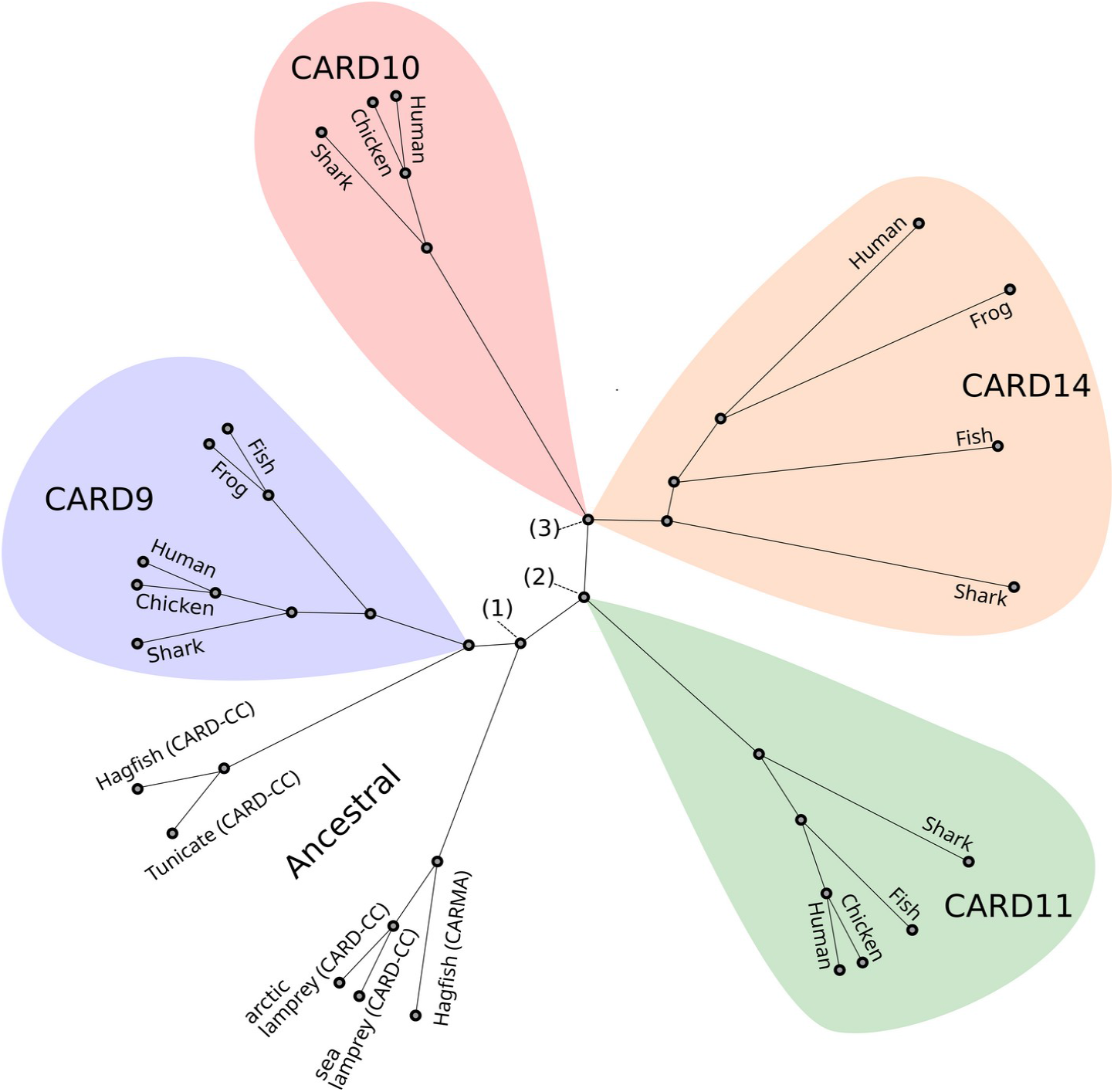
Relationship between the different CARD-CC family members. A phylogenetic (PhyML) tree from an alignment (MUSCLE) of vertebrate CARD-CC family members. As outgroup (root) and representative of the ancestral CARD-CC, sequences from two sequenced lamprey species (arctic: *Lethenteron camtschaticum* and sea: *Petromyzon marinus)*, hagfish (*Eptatretus burgeri*) and the invertebrate tunicate (*Ciona intestinalis*) were used. A likely evolutionary scenario is that **(1)** the ancestral CARD-CC gene got duplicated into CARD9 and the original ancestral CARMA (which got fused to a MAGUK domain from a Dlg-5/ZO like gene) before the last common vertebrate ancestor. **(2)** The ancestral CARMA then duplicated into CARD11 and the CARD-[10/14] ancestor. Finally **(3)**, the CARD-[10/14] ancestor got duplicated into CARD10 and CARD14. The events **(2-3)** occurred during vertebrate evolution after *agnathans* (e.g. lampreys and hagfishes) and before *chondrichthyes* (e.g. sharks and rays; “Shark”; *Callorhinchus milii*) branched off from the rest of the vertebrates (*euteleostomi*; e.g. bony fishes (“Fish”; *Danio rerio*), amphibians (“Frog”; *Xenopus tropicalis*), reptiles (“Chicken”; *Gallus gallus*) and mammals (“Human”; *Homo sapiens*)).

Receptor signaling via the ITAM (immunoreceptor tyrosine-based activation motif) binding one of the two related SH2 domain tyrosine kinases Zap70 or Syk activates PKC, leading to CARD-CC phosphorylation and the recruitment of a pre-existing Bcl10/MALT1 complex and TRAF6 (Fig. 2A) [11,13,33,34]. An alternative CBM-dependent pathway upstream of PKC activation is represented by the Gα_12_/Gα_13_ and RhoA-dependent G protein coupled receptors (GPCRs) [35]. MALT1 in the activated CBM complexes recruits critical downstream proteins, such as TRAF6, for activation of NF-κB-dependent gene expression (Fig. 2A) [36]. Apart from this NF-κB-activating “scaffold” function, MALT1 (PCASP1 [37]) also is a protease (paracaspase), which is a drugable catalytic activity. The anti-inflammatory role of many of the known MALT1 protease substrates coupled with the critical role for MALT1 in pro-inflammatory signaling has sparked an interest in targeting MALT1 protease activity as a therapeutic strategy for autoimmune diseases [38]. The proteolytic activity of MALT1 was also found to be critical for certain cancers [39], which has stimulated an interest in MALT1 protease activity as a cancer therapy target as well. The CBM complex signaling is a nice illustration of the double edged role of immune signaling components and the importance for their tight regulation [5]. For example, MALT1 in T cells is critical for immunity to an avirulent strain of Rabies virus [40], but at the same time is T cell-expressed MALT1 also contributing to mortality after infection by a virulent strain of Rabies virus due to excessive inflammation [41]. Disrupted MALT1 protease activity specifically in T cells is also leading to the development of spontaneous autoimmune disease [42].

**Fig. 2.**
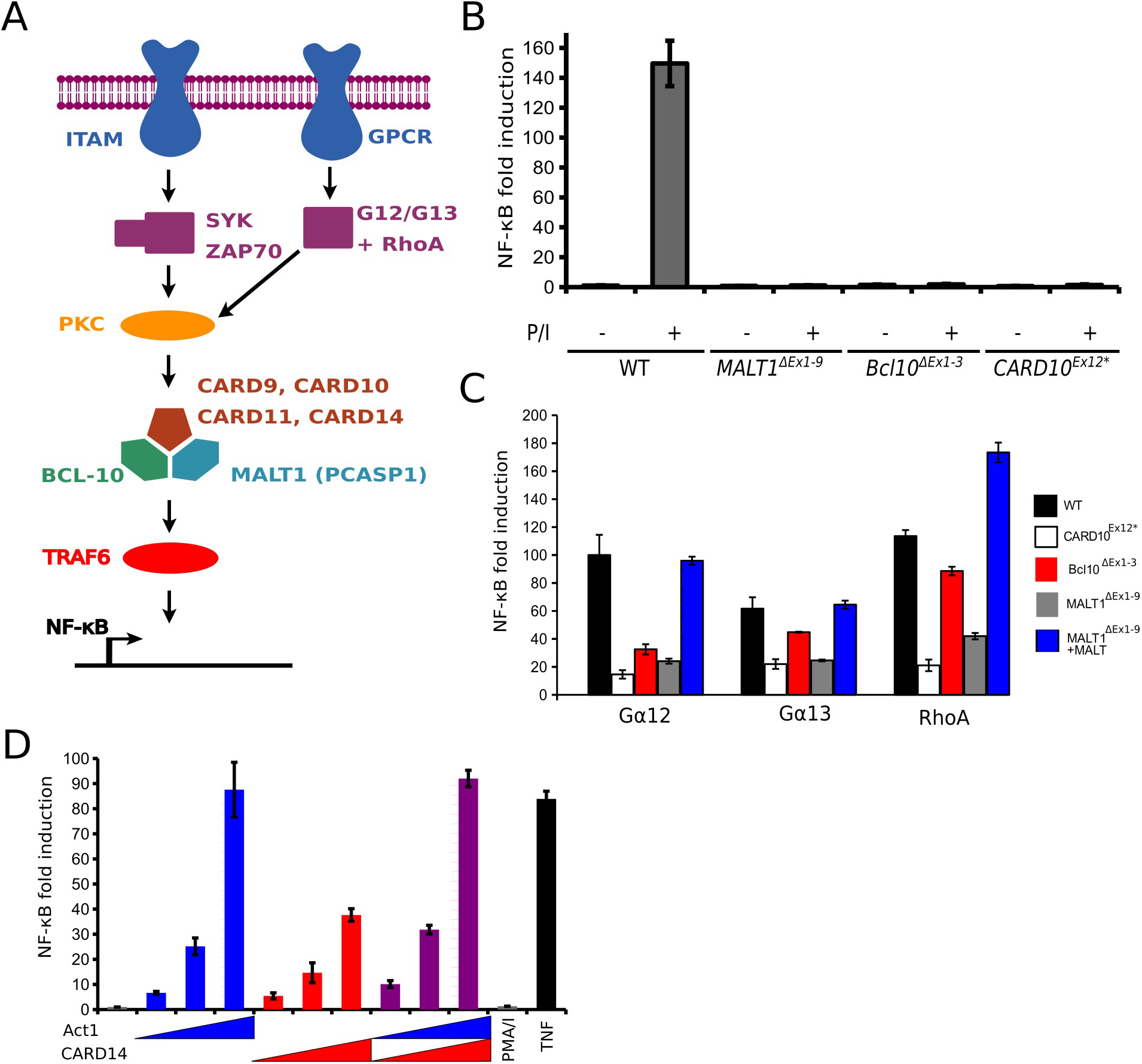
**A)** CBM complex signaling model. Upstream ITAM-containing receptors activate different types of PKC via the activity of the SH2 domain tyrosine kinases Syk or Zap70. Upon phosphorylation, the CBM complex is assembled, which recruits TRAF6 and activates downstream NF-κB transcriptional activity. **B)** PMA/Ionomycin (P/I)-induced NF-κB driven luciferase expression in wild-type (WT) and *MALT1, Bcl10* and *CARD10* mutant HEK293T cells after 6 hours stimulation. Fold-induction in PMA-stimulated (+) cells is compared to the average background level of the non-stimulated (-) cells. **C)** Gα_12_-Q231 L, Gα_13_-Q226L and RhoA-Q63L-induced NF-κB driven luciferase expression in wild-type (WT) and *MALT1, Bcl10* and *CARD10* mutant HEK293T cells. *MALT1* KO cells also reconstituted by transient expression of WT *MALT1*. D) To investigate the proposed Act1-CARD14 signaling axis, a gradient (0, 3, 10, 30 ng/μg transfected DNA) of Act1, CARD14 or Act1 and CARD14 together was evaluated in *CARD10* KO cells. As controls for the *CARD10* KO cells, PMA/ionomycin (P/I) and TNF stimulation were used. Luciferase values in **B-D** represent LacZ-normalized luciferase enzymatic activity expressed as fold induction (unitless). Error bars represent 95% confidence intervals (Student’s t distribution). Results are representative for three independent experiments.

Recently, a novel CARD-CC-independent oncogenic Bcl10/MALT1 activation mechanism has also been described, where the Karposi’s sarcoma-associated herpes virus (KSHV) virulence factors K15 and K13 engage directly with Bcl10/MALT1 in a “KBM” complex [43,44]. MALT1 can also be activated independently of Bcl10 and CARD-CC proteins, which is illustrated by that MALT1 homologs are present in species lacking Bcl10 and CARD-CC [11,45] and that MALT1 can be activated by TRAF6 overexpression in mammalian knock-out cells lacking Bcl 10 or CARD-CC proteins [46]. However, in the context of the dominant CBM complex-mediated MALT1 activation pathway, studies on CARD11 have shown that phosphorylation leads to release of intramolecular auto-inhibition and aggregation of the CARD-CC protein together with the other CBM complex components in huge macromolecular filamentous structures that acts as a platform for downstream signaling [47–49]. Many phosphorylation sites in CARD11 are highly conserved [5]. CARD9 has been shown to be activated by PKCδ [50], CARD10 is activated by PKCα and PKCθ [51,52] and CARD11 is activated by PKCβ and PKCθ in B and T cells, respectively [52–55]. It is very likely that there are more CBM-activating PKC isozymes, and that there are more PKC::CARD-CC kinase/substrate relationships in different cells and pathways. While the CARD-CC protein family is old and originated before the last common ancestor of planulozoa (cnidaria and bilateria) [11], the PKC family is however significantly older with origins in the last ancestor of opisthokonts (metazoa and fungi) [56], highlighting that PKCs also have many CBM-independent functions. In humans and mice, the PKC family consists of nine true members in the “conventional” (α, β, γ), “novel” (δ, ε, η, θ) and “atypical” (ζ, ι/λ) sub-classes [57,58]. The different PKCs show different substrate specificity [59,60], and have non-redundant biological functions [61]. The extended definition of the PKC superfamily also includes the related three PKN and three PRKD kinase family members [57]. PRKD1 was sometimes called PKCμ, and PRKD3 was called PKCν. Aberrant activation of PKC has been associated to several types of cancer [51,62,63], indicating that at least some of those might represent an oncogenic process dependent on over-active CBM signaling complexes. Some PKCs can however also act as tumor suppressors [61], which indicates that the involvement of this pathway is complex. The aim of this study is to discover the full scope of CBM complex dependent signaling pathways downstream of PKC activation. Defining the full scope of the functions of the CARD-CC/Bcl10/MALT1 complex-dependent pathways could ultimately be important in the context of future MALT1 inhibitor-based therapies [64,65].

## Results

### Verification of the screening tools for pathway analyses

A general activator of conventional and novel PKCs is the diacylglycerol (DAG) analog phorbol 12-myristate 13-acetate (PMA) [63]. PMA is often used together with Ionomycin due to its synergistic effects [66]. Ionomycin also provides the Ca^2+^ signal that is important for the activation of the conventional PKCs. To functionally verify that we have generated a cell line suitable for screening of PKC/CARD-CC interactions, we treated wild-type and mutant HEK293T cells with PMA/Ionomycin for 6 hours to activate a broad range of PKCs present endogenously in HEK293T cells and measured NF-κB – driven luciferase expression via the luciferase enzymatic activity. HEK293T express several conventional (α, β), novel (δ, ε, η, θ) and atypical (ı, ζ) PKC isoforms [18,67], which means that up to six PKCs are potentially activated by PMA/Ionomycin in HEK293T cells. Importantly, HEK293T genomes are highly variable and selection bottle necks might cause major changes to the cells [68], which means that the PKC expression profile might not be true for all subclones of this cell line. Despite this, HEK293T cells are chosen as model cell because they easily can be transiently transfected at a very high efficiency. Bcl10 and MALT1 are common for all CBM complexes and the knock-out cell lines are thus completely un-responsive to PMA stimulation (Fig. 2B). CARD10 is the dominant CARD-CC isoform (transcript per million (TPM): 11 vs 0.4 for *CARD9*, 1.0 for *CARD11* and 0.8 for *CARD14;* significance cut off >1.0 TPM) in HEK293T cells [18]. In agreement with this, our CARD10^Ex12^* KO HEK293T clone was as un-responsive to PMA stimulation as the *MALT1* and *Bcl10* KO clones (Fig. 2B). To evaluate the screening system and to confirm that the Gα_12_/Gα_13_-RhoA pathway [35] at least partially depend on CBM complex signaling, we transfected activated Gα_12_, Gα_13_ and RhoA mutants in the CARD10, Bcl10 and MALT1 KO HEK293T cells with or without MALT1 reconstitution (Fig. 2C). We used MALT1 KO cells for this pathway evaluation since reconstitution of MALT1 does not induce any NF-κB activity on its own, whereas overexpression of Bcl10 or CARD10 leads to spontaneous NF-κB activation. In contrast, another suggested upstream activator of a CBM complex-dependent signaling pathways is the IL-17 receptor adaptor Act1 [69]. In our screening system, Act1 activated NF-κB independently of the CBM complex components (Fig. 2D). Taken together, this demonstrates that KO HEK293T cells can be a useful tool for rapid evaluation of novel candidate upstream activation pathways.

### CARD-CC complementation and verification of activated PKC mutants

The dramatically reduced PMA responsiveness of the selected CARD10^Ex12^* mutant HEK293T clone (Fig. 2B) enables systematic analysis of specific PKC/CARD-CC interactions through reconstituton by transient transfection. CARD9 and CARD11 could easily be investigated using a default transfection set-up in the CARD10^Ex12^* mutant HEK293T cells since basal activation by overexpression was very low (Fig. 3A). CARD10 and CARD14, on the other hand, seem to have a much poorer auto-inhibition (or they get activated by cellular events triggered by calcium phosphate transfection), and in order to observe synergistic effects with activated PKCs, we had to titrate down the amount of expression plasmid DNA to the level where the basal activation was closer to that of CARD9 and CARD11 (Fig. 3A). Consistent with the poor auto-inhibition seen in the CARD10 and CARD14 transfections, further activation of CARD10 and −14 by the highly conserved L/LI insertion mutation was weak or not significant at high transfection levels (Fig. 3B), in contrast to the already reported dramatic activations of the L225LI (L232LI) CARD11 and L213LI CARD9 mutations [70,71]. However, both CARD10 and CARD14 show clear activation by the L/LI mutation at lower transfection levels (Fig. 3B), indicating that we can evaluate CARD10 and CARD14 activation by transfecting low ammounts of expression vector. Using the levels of transfection of the CARD-CC protein expression constructs that resulted in the lowest spontaneous activation while still showing highest potential for detection of upstream activation (Fig 3A,B), we transiently reconstituted the CARD10 KO HEK293T cells. The transiently reconstituted CARD10 KO HEK293T cells were stimulated with PMA/Ionomycin for 6 hours and analyzed for NF-κB-dependent luciferase induction. Unreconstituted cells did not respond to stimulation (Fig. 2B, 3C), and all four CARD-CC proteins were able to restore some of the responsiveness to PMA/Ionomycin stimulation, demonstrating that all four CARD-CC proteins are activated by PKCs (Fig. 3C).

**Fig. 3.**
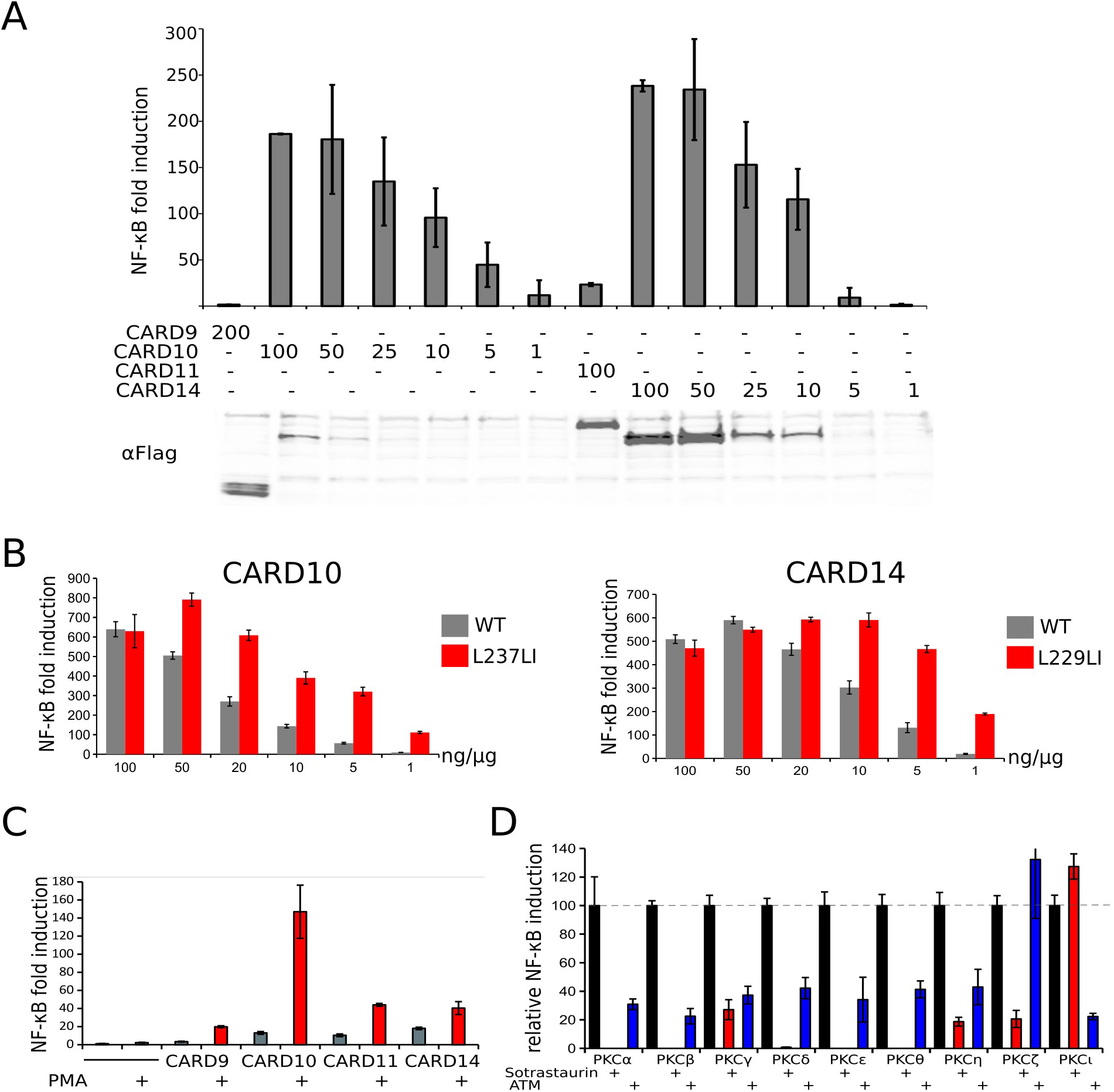
**A)** Basal NF-κB activation from CARD-CC overexpression vs empty vector transfected control, based on luciferase activity in wild-type HEK293T cells at different levels of transfection. CARD-CC expression determined by Western blot and anti-Flag tag antibody. **B)** Basal NF-κB activation in wild-type vs L/LI insertion mutants of CARD10 and CARD14 at different levels of transfection. For **A&B**, numbers indicate ng CARD-CC expression plasmid DNA/μg DNA master mix. **C)** PMA-induced NF-κB activation (red bars) after 6h stimulation in CARD10 KO HEK293T cells transiently reconstituted with the four different CARD-CC proteins. Grey bars represent NF-κB activation by expression of the CARD-CC proteins without stimulation. **D)** Relative NF-κB activation by activated PKC mutants in presence of PKC inhibitors. Uninhibited PKC (black bar, dashed line) is set to 100, red bars NF-κB activation in presence of Sotrastaurin and blue bars in presence of aurothiomalate (ATM). For Luciferase results in **A-D**, results are representative for four equivalent independent experiments and error bars represent 95% confidence intervals (Student’s t distribution).

Since PMA/Ionomycin most likely activates multiple PKCs in HEK293T cells, we made use of overexpression of activated PKC mutants in order to more specifically delineate the PKC/CARD-CC functional interactions. In order to verify that NF-κB activation from overexpression of the activated PKC mutants was due to their kinase activities, we first evaluated their NF-κB activating potential either non-treated or in presence of the PKC inhibitors Sotrastaurin or aurothiomalate (ATM) in wild-type HEK293T cells (Fig. 3D). The broad-range inhibitor Sotrastaurin could completely inhibit NF-κB-dependent luciferase induction downstream of most of the activated PKC isozymes. To see if the poor autoinhibition of CARD10 and CARD14 was due to a PKC signal induced by the calcium phosphate transfection, we also evaluated the basal activity of CARD10 and CARD14 overexpression in presence of two broad-range PKC inhibitors. Neither Gö6983 nor Sotrastaurin reduced the spontaneous activity of overexpressed CARD10 or CARD14, indicating that overexpression-induced activation is independent of upstream PKC signals (not shown).

In conclusion, since we could verify that the PKC-induced NF-κB activation was kinase activity and CBM complex dependent, and that we had managed to make a model cell line where the dominant CARD-CC was deleted in HEK293T cells, we are confident to use these tools for screenings of synergistic PKC/CARD-CC interactions.

### Identification PKC/CARD-CC functional interactions

Analysis of CARD9 activation by overexpression of activated PKCs revealed a surprisingly strong novel activation by PKC-η, −θ and −ε, along with the published PKCδ [50] (Fig. 4A). CARD10, on the other hand, gets activated by a wide range of PKCs (Fig. 4B), including the atypical sub-class, which was inactive against CARD9 (Fig. 4A). This demonstrates that different CARD-CC proteins get differentially activated by different PKCs. As expected, CARD11 gets activated by PKCβ and PKCθ but CARD11 is also activated by PKCε and PKCη (Fig. 4C), making the CARD11 activation profile more CARD9-like than CARD10-like. In contrast to the other CARD-CC proteins, no upstream receptor pathway is known for CARD14, where activated mutations are implicated in psoriasis [14]. The autoinhibition is less pronounced than for CARD9 and CARD11 (Fig. 3A). By careful adjustment of the CARD14 expression level, we could however verify that also CARD14 gets activated by PKCδ and to a much lesser extent by several other PKCs (Fig. 4D). Despite having approximately the same basal activation level as CARD10 (Fig. 3A), CARD14 responded much less to activated PKCs. There are however still some limitations in the screening system. For example, the activated PKCα was very poorly expressed (Fig. 4E) and in order to see any clear effects, it had to be transfected at very high levels. Because of this, the effects seen by this PKC could be under-estimated due to a limitation of the screening system using activated PKC mutants. It is known that some PKCs get destabilized and degraded upon activation [72]. Due to the massively different protein expression levels between the different activated PKC isoforms, protein expression from luciferase experiments from Fig. 4A-D were evaluated per PKC (Fig 4E). In conclusion, systematic evaluation of all pairwise PKC and CARD-CC functional interactions revealed a surprising number of highly potent synergistic activating interactions. It is however important to note that the very high fold inductions seen for CARD9 more has to do with a very low basal activation of CARD9 compared to the other CARD-CC proteins (Fig. 4A-D), which could be due to an additional layer of autoinhibition [73]. Similarly, some of the PKC contributions could very well be underestimated. The very poor protein expression of activated PKCα could mean that the CARD-CC activating potential of this particular isozyme is underestimated. Howevwer, as a counterexample is PKCε one of the strongest inducers of CARD-CC activity despite showing very poor protein expression (Fig. 4).

**Fig. 4.**
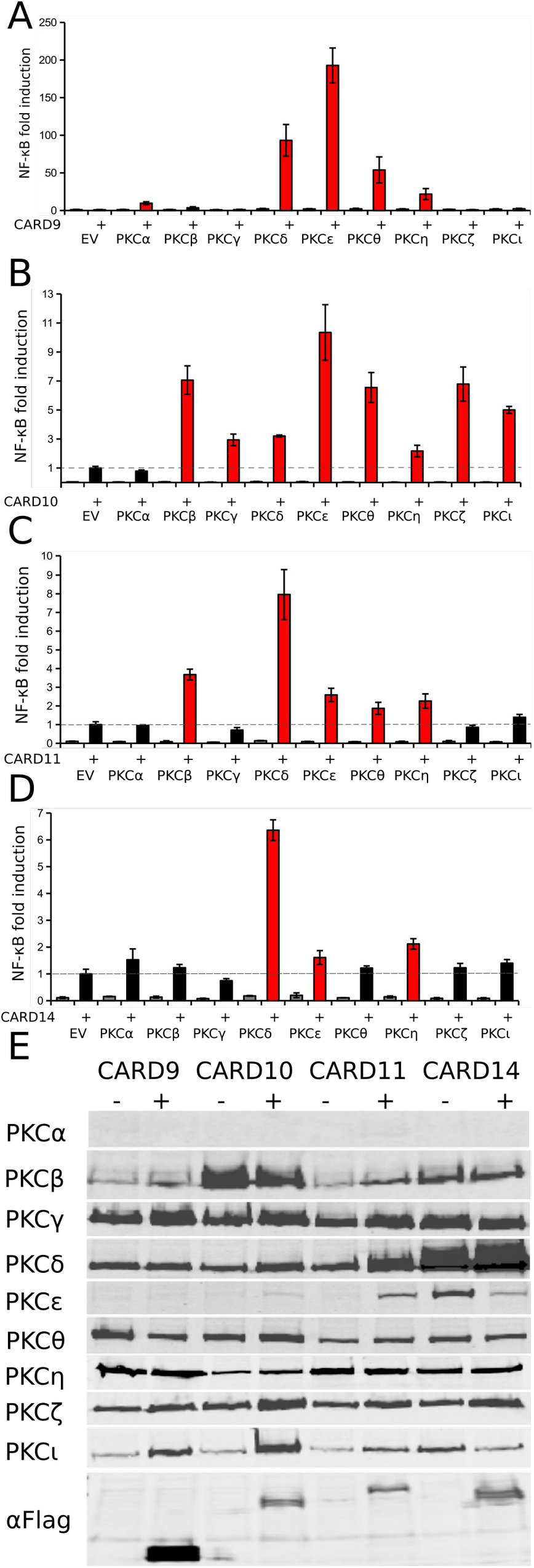
PKC-dependent NF-κB activation via **A)** CARD9, **B)** CARD10, **C)** CARD11, and D) CARD14. For **A-D**, activation by activated PKC mutants expressed as fold-induction of NF-κB-induced luciferase expression, compared to CARD-CC expressed without activated PKC (EV, set to 1, dashed line in **B-D**). Luciferase results are representative for four equivalent independent experiments and error bars represent 95% confidence intervals (Student’s t distribution). Red bars represent PKC/CARD-CC synergistic activation that is significantly (p<0.05) greater than the CARD-CC expressed alone. **E)** Expression of PKCs and CARD-CC proteins in experiments **A-D**.

### CARD-CC heterocomplex formation is a confounding factor in PKC response profiles

For reliable PKC::CARD-CC screenings, the isolation of a true CARD10 KO CARD10^Ex12^* clone was critical. Initial screenings were done in an incomplete CARD10 KO CARD10^Ex12^* HEK293T mutant (#C8) which showed approximately 10% NF-κB activation after PMA stimulation compared to WT cells. This kind of incomplete KO from frame shift mutation indels can occur by multiple mechanisms [74]. Two PKCs (PKCβ and PKCδ) show relatively high basal activity in WT cells. This high background was also present in the incomplete CARD10^Ex12^* #C8 mutant HEK293T clone without CARD-CC complementation, which caused an underestimation of their activating potential. Using the completely functional CARD10 KO CARD10^Ex12^* #C62 clone, NF-κB activation by those two PKCs were as negative as the other PKCs without CARD-CC complementation (Fig. 4A-D). For CARD9, −10 and −11 the general PKC response pattern was highly reproducible in experiments from the #C8 and #C62 clone, albeit with a relative under-estimation of the contribution from PKCβ and PKCδ in #C8. In contrast, CARD14 showed a dramatically different pattern between the incomplete #C8 and the complete #C62 clone: In presence of ~10% remaining CARD10 actvity, the PKC response pattern of CARD14 resembled that of CARD10, whereas a clean CARD10-deficient background revealed a highly specific CARD14 activation via PKCδ (Fig. 4D). This discrepancy might indicate that there is some kind of CARD10/CARD14 cross-talk in cells expressing both isoforms, such as developing keratinocytes [75]. We could also verify that presence of CARD10 led to a different CARD14 PKC response pattern by transient reconstitution in the complete *CARD10* KO clone #62, where the CARD10 effect appears dominant with only a minor contribution from CARD14 in presence of PKCδ or PKCη (Fig. 5A). To investigate this further, we evaluated the capacity of CARD14 to form heterocomplexes with the other CARD-CC family members by co-immunoprecipitation, which revealed that CARD14 can form heterocomplexes with CARD10 and CARD11 but not CARD9 (Fig. 5B).

**Fig. 5.**
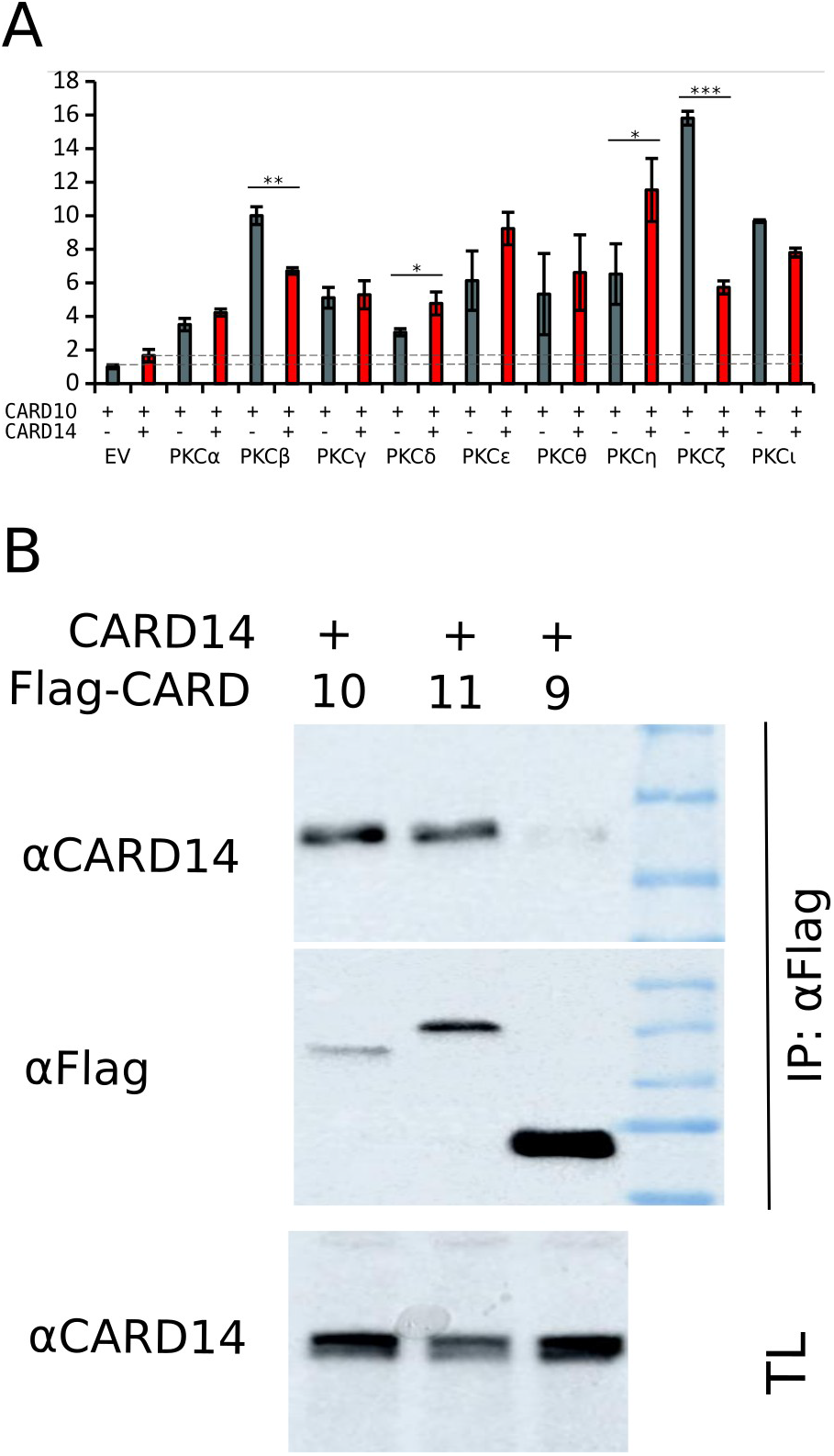
**A)** Investigation of the CARD14 PKC activation profile in presence of CARD10. Grey bars indicate CARD10+PKC activation profile and red bars CARD10+CARD14+PKC activation profile. **B)** Co-immunoprecipitation of un-tagged CARD14 with flag-tagged CARD9, −10 or −11.

### Evaluation of a common candidate site for PKC-mediated activation of CARD9 and CARD14

An interesting question is whether all activating PKCs are targeting the same phosphorylation sites on a given CARD-CC protein, or if CARD-CC activation by PKC occurs via several different phosphorylation events. CARD9 has been shown to be activated through phosphorylation at T231 by PKCδ [50], and PKCθ has been shown to activate CARD10 by phosphorylation on S520 and CARD11 by phosphorylation at S552/ S555 [52]. CARD11 is phosphorylated at similar sites by PKCβ [55], indicating that different PKCs activate CARD-CC proteins by phosphorylating common sites. One possible explanation for why CARD14 was unable to complement CARD11 deficiency in T cells [52] could be that CARD14 is not activated by PKCθ (Fig. 4D).

To investigate whether all PKCs could activate CARD9 via the same phosphorylation event, we compared the PKC response of WT and T231A mutant CARD9 (Fig. 6A-B). This clearly shows that for CARD9, all activating PKCs depend on the same residue for activation. And importantly, the T231A mutation does not affect the phosphorylation-independent overexpression-induced basal activity from CARD9 (Fig. 6A). The T231 site on CARD9 is also highly interesting in the context of CARD14 activation. Zymosan treatment on keratinocytes has been proposed to trigger an upstream signal of CARD14 [16], and zymosan responses signal via PKCδ to CARD9 in innate immune cells [50]. Interestingly, PKCδ is also the most potent activator of CARD14 (Fig. 4D). CARD14 S250 nicely aligns with CARD9 T231 and the oncogenic CARD11 S250P mutation [11], indicating that this is a common and evolutionarily conserved activation site. To evaluate whether CARD14 S250 also would be responsible for PKCδ-mediated activation of CARD14, we evaluated the CARD14 S250A mutant both for spontaneous activity (Fig. 6C) and for PKC responses (Fig. 6D). In contrast to the CARD9 T231A mutation, the CARD14 S250A mutation only partially reduced PKCδ-mediated activation of CARD14 and did not negatively influence activation by PKCε or PKCη, indicating that alternative sites contribute to CARD14 activation (Fig. 6D). This is the first time that it has been clearly demonstrated that also CARD14 gets activated by upstream PKCs, and it will be very interesting to identify the additional phosphorylation site(s) for PKC-mediated CARD14 activation.

**Fig. 6.**
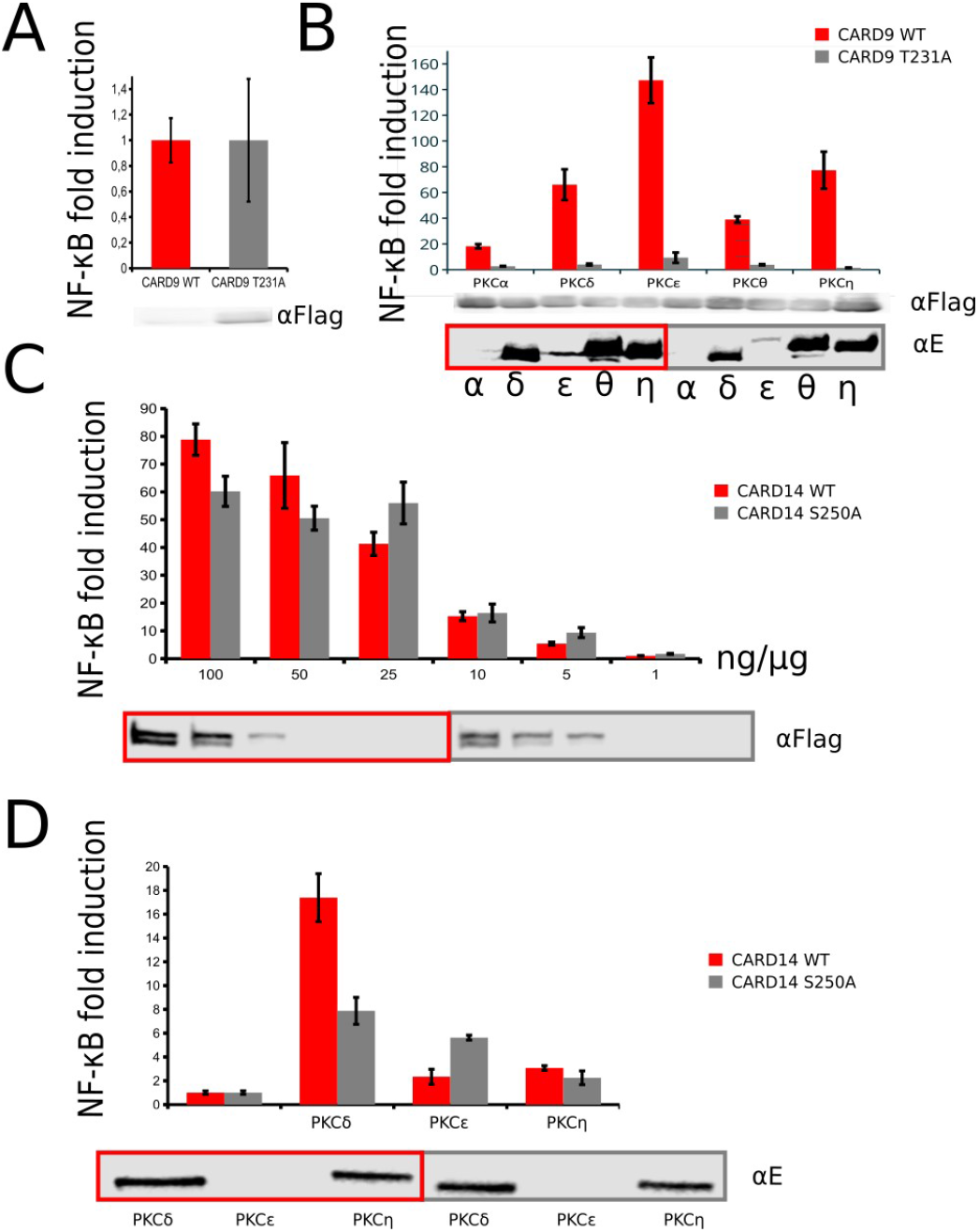
Different PKCs utilize the same critical phosphorylation site for CARD9 activation, but not for CARD14. **A)** Basal NF-κB activation by CARD9 and CARD9-T231A **B)** Activation of CARD9 and CARD9-T231A in presence of the CARD9-activating PKCs **C)** Levels of spontaneous NF-κB activation in a transfection gradient of CARD14 and CARD14-S250A. **D)** NF-κB by CARD14 and CARD14-S250A in presence of CARD14-activating PKCs.

## Discussion

Our novel screening system for functional interactions uncovered several previously unknown potential pathways which can be further explored in the future (Fig. 7). An interesting general conclusion is that the PKC response profiles do seem to correlate with the evolutionary relationships in the CARD-CC family (Fig. 1), where CARD9 and CARD11 have a very similar PKC response profile (Fig. 4) and CARD10 and CARD14 appear to be able to functionally interact (Fig. 5). This phylogenetic functional division might also expand to their expression profiles, where CARD9 and CARD11 are expressed in immune cells and CARD10 and CARD14 in non-hemapoetic cells. Given the potential correlation between phylogenetic relations and functional similarity, it would be highly interesting to also investigate the PKC response profile in ancestral CARD-CC by comparing related isoforms from organisms like jawless vertebrates and tunicates. Based on the phylogeny of the CARD-CC family (Fig. 1), we would expect those forms to have a CARD9/11-like activation profile. If the ancestral CARD-CC isoforms also show a CARD9/11-like restricted expression to immune-related cells (see for example: [76–78]), it would mean that the adoption of CBM complex activation downstream of other PKCs and in other cell types probably only evolved in the jawed vertebrate lineage after the expansion of the CARD-CC family. Supporting this hypothesis, the CARD-CC transcript (g47136) is most expressed in one “large lymphocyte”-like hemapoetic stem cell population (CP25) in the golden star tunicate [78]. The overlapping PKC response profiles of CARD9 and CARD11 are also interesting in the context of PKC functional evolution, since their typical upstream activators PKCδ and PKCθ share a relatively recent pre-vertebrate common ancestry [79].

**Fig. 7.**
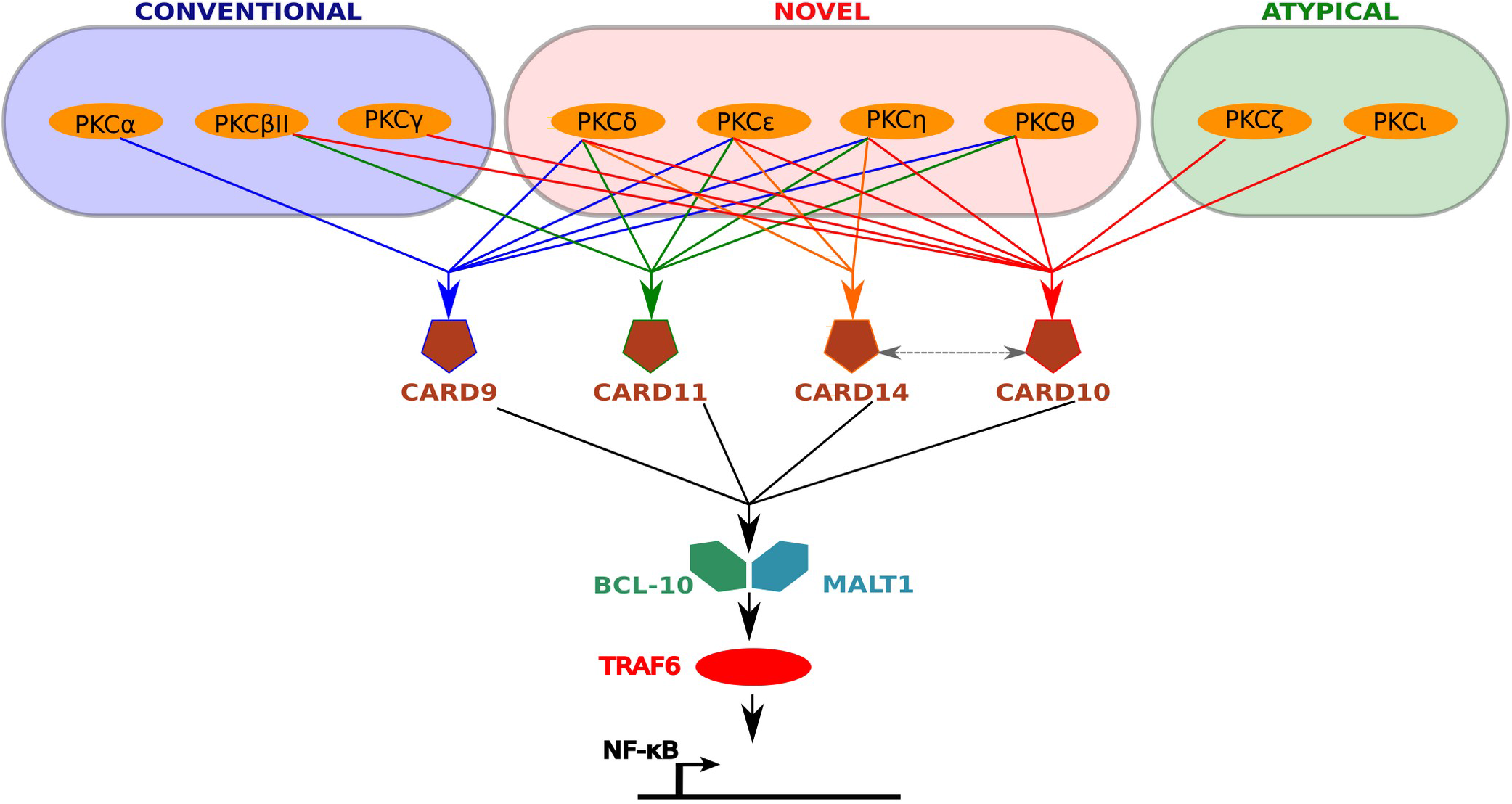
In conclusion, the different CARD-CC family members show different specificities towards upstream PKCs. All CARD-CC members are activated by the novel class of PKCs and only CARD10 can be activated by the atypical class of PKCs.

The different PKC response profiles of the different CARD-CC proteins also provides the support to test the involvement of CBM-complex dependent signaling in novel pathways. For example, some of the novel functional interactions uncovered by this study (like for example PKCη and CARD9) indicate interesting theoretical connections. It has been shown that PKCη is activated by PMA and other stimuli in myeloid cells leading to foam cell formation [80], which thus could be a CARD9 CBM complex-dependent process. Other generally CBM-activating PKC isoforms like PKCε are known to be involved in a wide range of inflammation- and cancer-related processes [81]. The specific PKC::CARD-CC interactions also opens up for a more complex and cell type-specific role for CBM-complex mediated signaling. For example is TNF generally considered to be a CBM-independent stimulus, mostly based on *in vitro* TNF stimulation of knock-out mouse embryonic fibroblasts (MEFs) [34]. TNF is however known to signal via PKCδ in neutrophils [82], and it is thus possible that TNF also activates CBM-dependent signaling pathways in some cell types but not others depending on their expression of specific PKC and CARD-CC isoforms. Since most cells express a complex mix of PKC isoforms, it is however challenging to study the specific *in vivo* functional interactions between one PKC and one CARD-CC isoform. However, given the specific requirement for T231 for CARD9 activation by multiple PKCs (Fig. 6B) and the different substrate recognition sequences for different PKCs [59,60], it might be possible to engineer CARD9 mutants that remain responsive to some PKCs but are insensitive to others. That way, one could establish the relative importance of different upstream PKC pathways for CARD9 function. Whether a similar strategy would work for the other CARD-CC family members requires further investigations. In contrast to the PKCs, which show a complex expression pattern across cell types, the CARD-CC family members are mostly restricted to specific and non-overlapping cell types [18,19]. Interesting in this context is our observation that transient complementation of *CARD10* KO HEK293T cells worked much better with CARD10 compared to all the other CARD-CC members (Fig. 3C), which could indicate that different cells express other yet unknown factors that influence CARD-CC activity. What our artificial overexpression system also did not address are the inducible subcellular translocations of PKC isoforms. These translocations can be critical. For example, disrupted PKC translocation in microgravity is severely inhibiting the activation of several different immune cells [83–85]. Theoretically, a disrupted CBM-complex dependent signaling downstream of PKC could also be partially responsible for other physiological effects of microgravity, like osteoporosis [86,87].

The surprising dependency of CBM complexes downstream of the atypical PKCs further expands the potential use of CBM-interfering interventions as a therapeutic strategy. For example, the PKCζ → NF-κB pathway has been found to be important in cardiac hypertrophy, prostate- and colorectal cancer [88–90]. PKCı/λ has also been shown to act as an oncogene in many different types of cancer [90,91]. This demonstrates that our findings can lead to new hypotheses to be tested in different biological systems which ultimately might lead to novel pharmacological treatment opportunities, for example by using MALT1 protease inhibitors [64]. Since we surprisingly could find synergistic activations between all PKCs (including the atypical) and CARD-CC proteins, it is logical to consider the broader context of the entire PKC superfamily (including the PRKD and PKN proteins) as potential CBM activators. However, our screenings with PKN and PRKD clones did not indicate any CBM-complex dependent signaling to NF-κB (not shown). However, there may also be other CARD-CC activating mechanisms from other classes of kinases or other protein-modifying enzymes that are currently unknown [5], as suggested for DNA damage-induced CARD10 CBM activation [92]. Recently, it was also reported that Dok3 negatively regulates CARD9 activation by recruitment of protein phosphatase 1 (PP1) [93]. A similar negative regulation of CARD11 has been described via PP2A [94]. Considering the overlapping PKC-dependent activation mechanisms of the four CARD-CC family members, it will be interesting to also investigate whether there is an overlap also in the mechanisms of negative regulation of CARD-CC activation.

## Methods

### CARD-CC and activated PKC expression plasmids

Plasmids of the cloned genes were deposited in the BCCM/GeneCorner plasmid collection along with detailed descriptions of cloning strategy and plasmid sequence (http://www.genecorner.ugent.be/). The *in silico* cloning, graphical vector map generation, phylogenetic- and sequence analysis was done in UGENE (http://ugene.net/) [95] or via the Linux command line (muscle [96], phyml [97]). Sequences used for the CARD-CC phylogenetic analysis (Fig. 1) were previously published as supplemental data [11] together with 2 new hagfish sequences (ENSEBUT00000026558.1 and ENSEBUT00000027027.1 from ENSEMBL [98]). Sequences and the raw tree file with distance values can be found in supplemental data. Expression plasmids with wild-type Flag-tagged CARD9 (LMBP 9609), CARD10 (LMBP 10187), CARD11 and CARD14 (LMBP 9863) were used together with expression plasmids with E-tagged or Flag-tagged activated mutant PKCα-A25E (LMBP 8931; LMBP 8926), PKCβII-A25E (LMBP 8918; LMBP 9040), PKCγ-A24E (LMBP 8919; LMBP 9041), PKCδ-RR144/145AA (LMBP 8924; LMBP 8521), PKCε-A159E (LMBP 8920; LMBP 9042). PKCθ-A148E (LMBP 8925; LMBP 9045), PKCη-A160E (LMBP 8922; LMBP 9044), PKCζ-A119E (LMBP 8921; LMBP 9043) and PKCı-A129E (LMBP 8923; LMBP 9046). For set-ups without PKC transfection, an equal amount of pCAGGS (LMBP 2453) empty vector (EV) was used. To test the extended PKC family, PKN1-ΔN (LMBP 10603), PKN2-ΔN (LMBP 10606), PKN3-ΔN (LMBP 10602), PRKD2 (LMBP 10605), PRKD3 (LMBP 10604) and PRKD1-S738E/S742E (Addgene: 10810) [99] were used. Investigations of spontaneous CARD10 and CARD14 activations compared to the highly conserved L/LI insertion site were performed with CARD10-L237LI (LMBP 10268) and CARD14-L229LI (LMBP 10267). The CARD-CC phosphorylation site mutants CARD9-T231A (LMBP 11266) and CARD14-S250A (LMBP 11488) were generated by PCR-based mutagenesis in the pICOz (LMBP 11103; 11224; 11223) minimal vector backbone [100] prior to re-cloning into the original expression vectors. For evaluation of GPCR components proposed to act upstream of CBM-complex dependent signaling, we used Gα_12_-Q231 L, Gα_13_-Q226L [101], and RhoA-Q63L (LMBP 04658) [102] expression plasmids. Also the IL-17 receptor adaptor Act1 (pEF-hCIKS) was evaluated as a potential upstream activator of CBM dependent signaling [69] by transient transfection in the KO model cells. For studies of CARD-CC heterocomplex formation with CARD14, a non-tagged CARD14 (LMBP 9863) clone was used [14].

### Genome-editing of human model cell lines

As positive controls, we generated large-deletion MALT1^ΔEx1-9^ (clone #M24) and Bcl10^ΔEx1-3^ (clone #B51) knock-out HEK293T cell lines. *MALT1* was knocked-out by co-transfecting two Cas9::guide constructs targeting exon 1 (LMBP: 10533) and exon 9 (LMBP: 10401). *Bcl10* was knocked-out by co-transfecting two Cas9::guide constructs targeting exon 1 (LMBP: 10402) and exon 3 (LMBP: 10403). *CARD10^Ex12^\** (clone #C62) was mutated by using a single Cas9::guide construct targeting exon 12 (LMBP: 10406). Early screenings were done with an incomplete *CARD10^Ex12^\** clone (clone #C8) which showed ~10% NF-κB activity compared to WT after PMA stimulation. Two days after transfection, single-cell sorting for GFP-positive (=successfully transfected) cells was performed on an ARIA II (BD Biosciences) fluorescence activated cell sorter (FACS) machine. Individual clones were evaluated for knock-out by PCR and functional assays. The CARD10^Ex12^* mutation was checked by PCR of the genomic region with the primers GT-hCARD10-Ex12-F: TCCCTGCCCAATCAGACTT and GT-hCARD10-Ex12-R: GCTTGTGACTTGAGATTTGAAGG. Sequencing of the amplified region was done with the nested primers Seq-hCARD10-Ex12-F: AATTAAGCCTGGCTGAGCA and Seq-hCARD10-Ex12-R: ATGATCAGCATTTCCCAGGT. Alignment of the complex indel chromatogram with a wild-type reference chromatogram was done using the TIDE calculator (https://tide-calculator.nki.nl/) [103].

### Cell culture, transfection and inhibitors

HEK293T cells were grown under standard conditions (DMEM, 10% FCS, 5% CO_2_, 37 ^o^C) and transfected with the calcium phosphate method [104]. PKC inhibition was done with three PKC inhibitors: 2μM Sotrastaurin (Selleckchem), 10μM Gö6983 (Tocris Bioscience) or 50μM aurothiomalate (Sigma-Aldrich). Inhibitors were added before transfection and when media was refreshed 6h after calcium phosphate transfection. For PMA/Ionomycin stimulation, 200ng/ml PMA (Sigma-Aldrich) and 1μM Ionomycin (Calbiochem) was added to the cells 6 hours before lysing the cells.

### Luciferase reporter gene assay and analysis

For evalutation of NF-κB induction, candidate inducing constructs were co-transfected with a NF-κB-dependent luciferase reporter expression plasmid (LMBP: 3249) [105] and an actin promoter-driven β-galactosidase (LacZ) expression plasmid (LMBP: 4341) [106] as transfection control. The CARD-CC/PKC screening transfecion set-ups were based on a 1 μg DNA master mix containing 100 ng luciferase reporter plasmid and 100 ng β-galactosidase expression plasmid. The amount of the different CARD-CC expression plasmids was determined individually based on basal spontaneous activation from overexpression: CARD9 (200 ng), CARD10 (5 ng), CARD11 (100 ng) and CARD14 (5 ng). The remaining DNA (600-795 ng) consisted of expression plasmid DNA for the activated PKC constructs or empty vector (EV). The 1 μg master mix was transfected over 4 24-well plate wells (4 · 10^4^ cells per well). The cells used for luciferase analysis were washed with PBS and lysed in luciferase lysis buffer (25mM Tris pH7.8, 2mM DTT, 2mM CDTA, 10% glycerol, 1% Triton X-100). For the colorimetric determination (at 595nm) of β-galactosidase activity, chlorophenol red-β-D-galactopyranoside (CPRG) (Roche diagnostics) was used as a substrate. Luciferase activity was measured by using beetle luciferin (Promega) as a substrate and luminescence was measured with the GloMax^®^ 96 Microplate Luminometer (Promega). Luciferase data was repeated at least 3 times with similar patterns. Data processing and calculation of 95% confidence intervals was done in Libreoffice (www.libreoffice.org) Calc (Student’s t-distribution [107,108]).

### Immunoprecipitation

HEK293T cells were seeded at 1 × 10^6^ cells on 90-mm Petri dishes and transfected with a total of 5 μg DNA. Twenty-four hours later, cells were lysed in MALT1 IP buffer (20 mM Tris-HCl pH 7.5, 137 mM NaCl, 1.5 mM MgCl2, and 1% Triton X-100) supplemented with protease and phosphatase inhibitors. Cell extracts were pre-cleared with 1 lg of total rabbit IgG (sc-2027, Santa Cruz) and 15 μl protein G-Sepharose beads (GE Healthcare, Buckinghamshire, UK). Proteins were precipitated with 1 μg of anti-Flag antibody (F-3165, Sigma) for 2 h at 4°C followed by the addition of 15 μl protein G-Sepharose beads for 1 h. The beads were washed four times with 1 ml lysis buffer and resuspended in 40 μl Laemmli buffer (0.1% 2-mercaptoethanol, 5ppm bromophenol blue, 10% Glycerol, 2% SDS, 63 mM Tris-HCl (pH 6.8)) [109]. Proteins were separated by SDS-PAGE and analyzed by Western blotting.

### Protein expression analysis

Transfected cells for expression control in luciferase experiments were lysed directly in Laemmli buffer. Protein concentration in total lysate control samples for immunoprecipitation were first determined by the Bradford method [110] prior to addition of Laemmli buffer. Protein lysates were run on a 10% PAGE gel and blotted on nitrocellulose. Expression of the CARD-CC proteins was determined with anti-Flag antibody (F-3165, Sigma). Expression of the E-tagged PKC clones with anti-E-tag antibody (ab66152, Abcam) and un-tagged CARD14 was detected with anti-CARD14 (HPA023388, Sigma).

## Author contributions

J.S. designed the project, performed experiments and wrote the manuscript. Y.D., M.H., M.K., I.S., D.V., I.A. and H.B. performed experiments, R. B. supervised the work.

## Acknowledgments

We would like to thank Dr. Corinne Rösselet and Prof. Dr. Wim Declercq, Ghent University, for gateway entry vectors with activated mutant PKCs [111]. pSpCas9(BB)-2A-GFP (PX458) was a gift from Feng Zhang (Addgene plasmid # 48138) [112]. Flag-tagged CARD11 expression construct was a gift from Dr. Mathijs Baens, KU Leuven [113]. Gα_12_-Q231 L and Gα_13_-Q226L expression constructs were a kind gift from Prof. Dr. Yung Hou Wong (The Hong Kong University of Science and Technology, Hong Kong, China) [101]. Cell sorting was performed by Gert Van Isterdael, IRC, VIB-UGhent.

## Abbreviations

Bcl10: B Cell CLL/Lymphoma 10
CARD: Caspase activation and recruitment domain
CC: Coiled-coil domain
MALT1: Mucosa-associated lymphoid tissue lymphoma translocation protein 1
PKC: protein kinase C

## References

1 Afonina IS, Zhong Z, Karin M & Beyaert R (2017) Limiting inflammation-the negative regulation of NF-κB and the NLRP3 inflammasome. Nat. Immunol. 18, 861–869.

2 Hathcock D, Sheehy J, Weisenberger C, Ilker E & Hinczewski M (2016) Noise Filtering and Prediction in Biological Signaling Networks. IEEE Trans. Mol. Biol. Multi-Scale Commun. 2, 16–30.

3 Aminian G, Farahnak-Ghazani M, Mirmohseni M, Nasiri-Kenari M & Fekri F (2015) On the Capacity of Point-to-Point and Multiple-Access Molecular Communications With Ligand-Receptors. IEEE Trans. Mol. Biol. Multi-Scale Commun. 1, 331–346.

4 Gohari A, Mirmohseni M & Nasiri-Kenari M (2016) Information Theory of Molecular Communication: Directions and Challenges. IEEE Trans. Mol. Biol. Multi-Scale Commun. 2, 120–142.

5 Lork M, Staal J & Beyaert R (2019) Ubiquitination and phosphorylation of the CARD11-BCL10-MALT1 signalosome in T cells. Cell. Immunol. 340, 103877.

6 Friedlander T, Mayo AE, Tlusty T & Alon U (2015) Evolution of Bow-Tie Architectures in Biology. PLOS Comput. Biol. 11, e1004055.

7 Supper J, Spangenberg L, Planatscher H, Dräger A, Schröder A & Zell A (2009) BowTieBuilder: modeling signal transduction pathways. BMC Syst. Biol. 3, 67.

8 Tieri P, Grignolio A, Zaikin A, Mishto M, Remondini D, Castellani GC & Franceschi C (2010) Network, degeneracy and bow tie. Integrating paradigms and architectures to grasp the complexity of the immune system. Theor. Biol. Med. Model. 7, 32.

9 Iglesias PA (2016) The Use of Rate Distortion Theory to Evaluate Biological Signaling Pathways. IEEE Trans. Mol. Biol. Multi-Scale Commun. 2, 31–39.

10 Fang Y, Noel A, Yang N, Eckford AW & Kennedy RA (2017) Convex Optimization of Distributed Cooperative Detection in Multi-Receiver Molecular Communication. IEEE Trans. Mol. Biol. Multi-Scale Commun. 3, 166–182.

11 Staal J, Driege Y, Haegman M, Borghi A, Hulpiau P, Lievens L, Gul IS, Sundararaman S, Gonçalves A, Dhondt I, Pinzón JH, Braeckman BP, Technau U, Saeys Y, van Roy F & Beyaert R (2018) Ancient Origin of the CARD–Coiled Coil/Bcl10/MALT1-Like Paracaspase Signaling Complex Indicates Unknown Critical Functions. Front. Immunol. 9.

12 Gross O, Gewies A, Finger K, Schäfer M, Sparwasser T, Peschel C, Förster I & Ruland J (2006) Card9 controls a non-TLR signalling pathway for innate anti-fungal immunity. Nature 442, 651–6.

13 Che T, You Y, Wang D, Tanner MJ, Dixit VM & Lin X (2004) MALT1/paracaspase is a signaling component downstream of CARMA1 and mediates T cell receptor-induced NF-kappaB activation. J. Biol. Chem. 279, 15870–15876.

14 Afonina IS, Van Nuffel E, Baudelet G, Driege Y, Kreike M, Staal J & Beyaert R (2016) The paracaspase MALT1 mediates CARD14-induced signaling in keratinocytes. EMBO Rep. 17, 914–927.

15 Howes A, O’Sullivan PA, Breyer F, Ghose A, Cao L, Krappmann D, Bowcock AM & Ley SC (2016) Psoriasis mutations disrupt CARD14 autoinhibition promoting BCL10-MALT1-dependent NF-κB activation. Biochem. J., BCJ20160270.

16 Schmitt A, Grondona P, Maier T, Brändle M, Schönfeld C, Jäger G, Kosnopfel C, Eberle FC, Schittek B, Schulze-Osthoff K, Yazdi AS & Hailfinger S (2016) MALT1 Protease Activity Controls the Expression of Inflammatory Genes in Keratinocytes upon Zymosan Stimulation. J. Invest. Dermatol. 136, 788–797.

17 McAllister-Lucas LM, Ruland J, Siu K, Jin X, Gu S, Kim DSL, Kuffa P, Kohrt D, Mak TW, Nuñez G & Lucas PC (2007) CARMA3/Bcl10/MALT1-dependent NF-kappaB activation mediates angiotensin II-responsive inflammatory signaling in nonimmune cells. Proc Natl Acad Sci U A 104, 139–44.

18 Uhlen M, Oksvold P, Fagerberg L, Lundberg E, Jonasson K, Forsberg M, Zwahlen M, Kampf C, Wester K, Hober S, Wernerus H, Björling L & Ponten F (2010) Towards a knowledge-based Human Protein Atlas. Nat. Biotechnol. 28, 1248–1250.

19 Single-cell transcriptomics of 20 mouse organs creates a Tabula Muris (2018) Nature 562, 367.

20 Stepensky P, Keller B, Buchta M, Kienzler A-K, Elpeleg O, Somech R, Cohen S, Shachar I, Miosge LA, Schlesier M, Fuchs I, Enders A, Eibel H, Grimbacher B & Warnatz K (2013) Deficiency of caspase recruitment domain family, member 11 (CARD11), causes profound combined immunodeficiency in human subjects. J. Allergy Clin. Immunol. 131, 477–485.e1.

21 Compagno M, Lim WK, Grunn A, Nandula SV, Brahmachary M, Shen Q, Bertoni F, Ponzoni M, Scandurra M, Califano A, Bhagat G, Chadburn A, Dalla-Favera R & Pasqualucci L (2009) Mutations of multiple genes cause deregulation of NF-kappaB in diffuse large B-cell lymphoma. Nature 459, 717–721.

22 Lu HY, Bauman BM, Arjunaraja S, Dorjbal B, Milner JD, Snow AL & Turvey SE (2018) The CBM-opathies-A Rapidly Expanding Spectrum of Human Inborn Errors of Immunity Caused by Mutations in the CARD11-BCL10-MALT1 Complex. Front. Immunol. 9, 2078.

23 Brohl AS, Stinson JR, Su HC, Badgett T, Jennings CD, Sukumar G, Sindiri S, Wang W, Kardava L, Moir S, Dalgard CL, Moscow JA, Khan J & Snow AL (2015) Germline CARD11 Mutation in a Patient with Severe Congenital B Cell Lymphocytosis. J. Clin. Immunol. 35, 32–46.

24 Jordan CT, Cao L, Roberson EDO, Pierson KC, Yang C-F, Joyce CE, Ryan C, Duan S, Helms CA, Liu Y, Chen Y, McBride AA, Hwu W-L, Wu J-Y, Chen Y-T, Menter A, Goldbach-Mansky R, Lowes MA & Bowcock AM (2012) PSORS2 is due to mutations in CARD14. Am. J. Hum. Genet. 90, 784–795.

25 Oudes AJ, Roach JC, Walashek LS, Eichner LJ, True LD, Vessella RL & Liu AY (2005) Application of affymetrix array and massively parallel signature sequencing for identification of genes involved in prostate cancer progression. BMC Cancer 5, 86.

26 Peled A, Sarig O, Sun G, Samuelov L, Ma CA, Zhang Y, Dimaggio T, Nelson CG, Stone KD, Freeman AF, Malki L, Vidal LS, Chamarthy LM, Briskin V, Mohamad J, Pavlovsky M, Walter JE, Milner JD & Sprecher E (2019) Loss-of-function mutations in caspase recruitment domain-containing protein 14 (CARD14) are associated with a severe variant of atopic dermatitis. J. Allergy Clin. Immunol. 143, 173–181.e10.

27 Demeyer A, Van Nuffel E, Baudelet G, Driege Y, Kreike M, Muyllaert D, Staal J & Beyaert R (2019) MALT1-deficient mice develop atopic-like dermatitis upon aging. Front. Immunol. 10.

28 Zhou T, Souzeau E, Sharma S, Siggs OM, Goldberg I, Healey PR, Graham S, Hewitt AW, Mackey DA, Casson RJ, Landers J, Mills R, Ellis J, Leo P, Brown MA, MacGregor S, Burdon KP & Craig JE (2016) Rare variants in optic disc area gene CARD10 enriched in primary open-angle glaucoma. Mol. Genet. Genomic Med. 4, 624–633.

29 Nho K, Corneveaux JJ, Kim S, Lin H, Risacher SL, Shen L, Swaminathan S, Ramanan VK, Liu Y, Foroud T, Inlow MH, Siniard AL, Reiman RA, Aisen PS, Petersen RC, Green RC, Jack CR, Weiner MW, Baldwin CT, Lunetta K, Farrer LA, Multi-Institutional Research on Alzheimer Genetic Epidemiology (MIRAGE) Study, Furney SJ, Lovestone S, Simmons A, Mecocci P, Vellas B, Tsolaki M, Kloszewska I, Soininen H, AddNeuroMed Consortium, McDonald BC, Farlow MR, Ghetti B, Indiana Memory and Aging Study, Huentelman MJ, Saykin AJ & Alzheimer’s Disease Neuroimaging Initiative (ADNI) (2013) Whole-exome sequencing and imaging genetics identify functional variants for rate of change in hippocampal volume in mild cognitive impairment. Mol. Psychiatry 18, 781–787.

30 Grabiner BC, Blonska M, Lin P-C, You Y, Wang D, Sun J, Darnay BG, Dong C & Lin X (2007) CARMA3 deficiency abrogates G protein-coupled receptor-induced NF-{kappa}B activation. Genes Dev. 21, 984–996.

31 Scudiero I, Vito P & Stilo R (2014) The three CARMA sisters: so different, so similar: a portrait of the three CARMA proteins and their involvement in human disorders. J. Cell. Physiol. 229, 990–997.

32 McAuley JR, Freeman TJ, Ekambaram P, Lucas PC & McAllister-Lucas LM (2018) CARMA3 Is a Critical Mediator of G Protein-Coupled Receptor and Receptor Tyrosine Kinase-Driven Solid Tumor Pathogenesis. Front. Immunol. 9.

33 Ruefli-Brasse AA, French DM & Dixit VM (2003) Regulation of NF-kappaB-dependent lymphocyte activation and development by paracaspase. Science 302, 1581–4.

34 Ruland J, Duncan GS, Wakeham A & Mak TW (2003) Differential requirement for Malt1 in T and B cell antigen receptor signaling. Immunity 19, 749–58.

35 Yu OM & Brown JH (2015) G Protein-Coupled Receptor and RhoA-Stimulated Transcriptional Responses: Links to Inflammation, Differentiation, and Cell Proliferation. Mol. Pharmacol. 88, 171–180.

36 Sun L, Deng L, Ea C-K, Xia Z-P & Chen ZJ (2004) The TRAF6 ubiquitin ligase and TAK1 kinase mediate IKK activation by BCL10 and MALT1 in T lymphocytes. Mol Cell 14, 289–301.

37 Hulpiau P, Driege Y, Staal J & Beyaert R (2016) MALT1 is not alone after all: identification of novel paracaspases. Cell. Mol. Life Sci. CMLS 73, 1103–1116.

38 Mc Guire C, Elton L, Wieghofer P, Staal J, Voet S, Demeyer A, Nagel D, Krappmann D, Prinz M, Beyaert R & van Loo G (2014) Pharmacological inhibition of MALT1 protease activity protects mice in a mouse model of multiple sclerosis. J. Neuroinflammation 11, 124.

39 Juilland M & Thome M (2016) Role of the CARMA1/BCL10/MALT1 complex in lymphoid malignancies. Curr. Opin. Hematol. 23, 402–409.

40 Kip E, Staal J, Verstrepen L, Tima HG, Terryn S, Romano M, Lemeire K, Suin V, Hamouda A, Kalai M, Beyaert R & Gucht SV (2018) MALT1 controls attenuated rabies virus by inducing early inflammation and T cell activation in the brain. J. Virol., JVI.02029–17.

41 Kip E, Staal J, Tima HG, Verstrepen L, Romano M, Lemeire K, Suin V, Hamouda A, Baens M, Libert C, Kalai M, Van Gucht S & Beyaert R (2018) Inhibition of MALT1 decreases neuroinflammation and pathogenicity of virulent rabies virus in mice. J. Virol.

42 Demeyer A, Skordos I, Driege Y, Kreike M, Hochepied T, Baens M, Staal J & Beyaert R (2019) MALT1 Proteolytic Activity Suppresses Autoimmunity in a T Cell Intrinsic Manner. Front. Immunol. 10.

43 Juilland M, Bonsignore L & Thome M (2017) MALT1 protease activity in primary effusion lymphoma. Oncotarget 9, 12542–12543.

44 Bonsignore L, Passelli K, Pelzer C, Perroud M, Konrad A, Thurau M, Stürzl M, Dai L, Trillo-Tinoco J, Del Valle L, Qin Z & Thome M (2017) A role for MALT1 activity in Kaposi’s sarcoma-associated herpes virus latency and growth of primary effusion lymphoma. Leukemia 31, 614–624.

45 Flynn SM, Chen C, Artan M, Barratt S, Crisp A, Nelson GM, Peak-Chew S-Y, Begum F, Skehel M & Bono M de (2019) MALT1 mediates IL-17 Neural Signaling to regulate C. elegans behavior, immunity and longevity. bioRxiv, 658617.

46 Bardet M, Seeholzer T, Unterreiner A, Woods S, Krappmann D & Bornancin F (2018) MALT1 activation by TRAF6 needs neither BCL10 nor CARD11. Biochem. Biophys. Res. Commun. 506, 48–52.

47 Qiao Q, Yang C, Zheng C, Fontán L, David L, Yu X, Bracken C, Rosen M, Melnick A, Egelman EH & Wu H (2013) Structural Architecture of the CARMA1/Bcl10/MALT1 Signalosome: Nucleation-Induced Filamentous Assembly. Mol. Cell 51, 766–779.

48 David L, Li Y, Ma J, Garner E, Zhang X & Wu H (2018) Assembly mechanism of the CARMA1-BCL10-MALT1-TRAF6 signalosome. Proc. Natl. Acad. Sci. U. S. A. 115, 1499–1504.

49 Schlauderer F, Seeholzer T, Desfosses A, Gehring T, Strauss M, Hopfner K-P, Gutsche I, Krappmann D & Lammens K (2018) Molecular architecture and regulation of BCL10-MALT1 filaments. Nat. Commun. 9, 4041.

50 Strasser D, Neumann K, Bergmann H, Marakalala MJ, Guler R, Rojowska A, Hopfner K-P, Brombacher F, Urlaub H, Baier G, Brown GD, Leitges M & Ruland J (2012) Syk kinase-coupled C-type lectin receptors engage protein kinase C-σ to elicit Card9 adaptor-mediated innate immunity. Immunity 36, 32–42.

51 Mahanivong C, Chen HM, Yee SW, Pan ZK, Dong Z & Huang S (2008) Protein kinase Cα-CARMA3 signaling axis links Ras to NF-κB for lysophosphatidic acid-induced urokinase plasminogen activator expression in ovarian cancer cells. Oncogene 27, 1273–1280.

52 Matsumoto R, Wang D, Blonska M, Li H, Kobayashi M, Pappu B, Chen Y, Wang D & Lin X (2005) Phosphorylation of CARMA1 plays a critical role in T Cell receptor-mediated NF-kappaB activation. Immunity 23, 575–585.

53 Thuille N, Wachowicz K, Hermann-Kleiter N, Kaminski S, Fresser F, Lutz-Nicoladoni C, Leitges M, Thome M, Massoumi R & Baier G (2013) PKCθ/β and CYLD Are Antagonistic Partners in the NFκB and NFAT Transactivation Pathways in Primary Mouse CD3+ T Lymphocytes. PLOS ONE 8, e53709.

54 Shinohara H, Yasuda T, Aiba Y, Sanjo H, Hamadate M, Watarai H, Sakurai H & Kurosaki T (2005) PKC beta regulates BCR-mediated IKK activation by facilitating the interaction between TAK1 and CARMA1. J. Exp. Med. 202, 1423–1431.

55 Sommer K, Guo B, Pomerantz JL, Bandaranayake AD, Moreno-García ME, Ovechkina YL & Rawlings DJ (2005) Phosphorylation of the CARMA1 linker controls NF-kappaB activation. Immunity 23, 561–574.

56 Heinisch JJ & Rodicio R (2018) Protein kinase C in fungi—more than just cell wall integrity. FEMS Microbiol. Rev. 42.

57 Mellor H & Parker PJ (1998) The extended protein kinase C superfamily. Biochem. J. 332, 281–292.

58 Parker PJ & Murray-Rust J (2004) PKC at a glance. J. Cell Sci. 117, 131–132.

59 Nishikawa K, Toker A, Johannes F-J, Songyang Z & Cantley LC (1997) Determination of the Specific Substrate Sequence Motifs of Protein Kinase C Isozymes. J. Biol. Chem. 272, 952–960.

60 Kang J-H, Toita R, Kim CW & Katayama Y (2012) Protein kinase C (PKC) isozyme-specific substrates and their design. Biotechnol. Adv. 30, 1662–1672.

61 Newton AC (2018) Protein Kinase C: Perfectly Balanced. Crit. Rev. Biochem. Mol. Biol. 53, 208–230.

62 Totoń E, Ignatowicz E, Skrzeczkowska K & Rybczyńska M (2011) Protein kinase Cε as a cancer marker and target for anticancer therapy. Pharmacol. Rep. PR 63, 19–29.

63 Castagna M, Takai Y, Kaibuchi K, Sano K, Kikkawa U & Nishizuka Y (1982) Direct activation of calcium-activated, phospholipid-dependent protein kinase by tumor-promoting phorbol esters. J. Biol. Chem. 257, 7847–7851.

64 Demeyer A, Staal J & Beyaert R (2016) Targeting MALT1 Proteolytic Activity in Immunity, Inflammation and Disease: Good or Bad? Trends Mol. Med. 22, 135–150.

65 Meloni L, Verstrepen L, Kreike M, Staal J, Driege Y, Afonina IS & Beyaert R (2018) Mepazine Inhibits RANK-Induced Osteoclastogenesis Independent of Its MALT1 Inhibitory Function. Mol. Basel Switz. 23.

66 Chatila T, Silverman L, Miller R & Geha R (1989) Mechanisms of T cell activation by the calcium ionophore ionomycin. J. Immunol. Baltim. Md 1950 143, 1283–1289.

67 Atwood BK, Lopez J, Wager-Miller J, Mackie K & Straiker A (2011) Expression of G protein-coupled receptors and related proteins in HEK293, AtT20, BV2, and N18 cell lines as revealed by microarray analysis. BMC Genomics 12, 14.

68 Lin Y-C, Boone M, Meuris L, Lemmens I, Van Roy N, Soete A, Reumers J, Moisse M, Plaisance S, Drmanac R, Chen J, Speleman F, Lambrechts D, Van de Peer Y, Tavernier J & Callewaert N (2014) Genome dynamics of the human embryonic kidney 293 lineage in response to cell biology manipulations. Nat. Commun. 5, 4767.

69 Wang M, Zhang S, Zheng G, Huang J, Songyang Z, Zhao X & Lin X (2018) Gain-of-Function Mutation of Card14 Leads to Spontaneous Psoriasis-like Skin Inflammation through Enhanced Keratinocyte Response to IL-17A. Immunity 49, 66–79.e5.

70 De Bruyne M, Hoste L, Bogaert DJ, Van den Bossche L, Tavernier SJ, Parthoens E, Migaud M, Konopnicki D, Yombi JC, Lambrecht BN, van Daele S, Alves de Medeiros AK, Brochez L, Beyaert R, De Baere E, Puel A, Casanova J-L, Goffard J-C, Savvides SN, Haerynck F, Staal J & Dullaers M (2018) A CARD9 founder mutation disrupts NF-κB signaling by inhibiting BCL10 and MALT1 recruitment and signalosome formation. Front. Immunol. 9.

71 Lamason RL, McCully RR, Lew SM & Pomerantz JL (2010) Oncogenic CARD11 mutations induce hyperactive signaling by disrupting autoinhibition by the PKC-responsive inhibitory domain. Biochemistry 49, 8240–8250.

72 Lu Z, Liu D, Hornia A, Devonish W, Pagano M & Foster DA (1998) Activation of Protein Kinase C Triggers Its Ubiquitination and Degradation. Mol. Cell. Biol. 18, 839–845.

73 Holliday MJ, Witt A, Rodríguez Gama A, Walters BT, Arthur CP, Halfmann R, Rohou A, Dueber EC & Fairbrother WJ (2019) Structures of autoinhibited and polymerized forms of CARD9 reveal mechanisms of CARD9 and CARD11 activation. Nat. Commun. 10, 3070.

74 Tuladhar R, Yeu Y, Piazza JT, Tan Z, Clemenceau JR, Wu X, Barrett Q, Herbert J, Mathews DH, Kim J, Hwang TH & Lum L (2019) CRISPR-Cas9-based mutagenesis frequently provokes on-target mRNA misregulation. Nat. Commun. 10, 1–10.

75 Israёl L, Bardet M, Huppertz A, Mercado N, Ginster S, Unterreiner A, Schlierf A, Goetschy JF, Zerwes H-G, Roth L, Kolbinger F & Bornancin F (2018) A CARD10-Dependent Tonic Signalosome Activates MALT1 Paracaspase and Regulates IL-17/TNF-α-Driven Keratinocyte Inflammation. J. Invest. Dermatol. 138, 2075–2079.

76 Flajnik MF (2014) Reevaluation of the Immunological Big Bang: comparisons of two vertebrate adaptive immune systems. Curr. Biol. CB 24, R1060–R1065.

77 Franchi N & Ballarin L (2017) Immunity in Protochordates: The Tunicate Perspective. Front. Immunol. 8.

78 Rosental B, Kowarsky M, Seita J, Corey DM, Ishizuka KJ, Palmeri KJ, Chen S-Y, Sinha R, Okamoto J, Mantalas G, Manni L, Raveh T, Clarke DN, Tsai JM, Newman AM, Neff NF, Nolan GP, Quake SR, Weissman IL & Voskoboynik A (2018) Complex Mammalian-like Hematopoietic System Found in a Colonial Chordate. Nature 564, 425–429.

79 Chen Z, Gong B-N, Wang Q, Xiao Z, Deng C, Wang W & Li Y (2019) Characterisation of amphioxus protein kinase C-δ/θ reveals a unique proto-V3 domain suggesting an evolutionary mechanism for PKC-θ unique V3. Fish Shellfish Immunol. 84, 1100–1107.

80 Lee H-K, Yeo S, Kim J-S, Lee J-G, Bae Y-S, Lee C & Baek S-H (2010) Protein kinase C-eta and phospholipase D2 pathway regulates foam cell formation via regulator of G protein signaling 2. Mol. Pharmacol. 78, 478–485.

81 Capuani B, Pacifici F, Pastore D, Palmirotta R, Donadel G, Arriga R, Bellia A, Di Daniele N, Rogliani P, Abete P, Sbraccia P, Guadagni F, Lauro D & Della-Morte D (2016) The role of epsilon PKC in acute and chronic diseases: Possible pharmacological implications of its modulators. Pharmacol. Res. 111, 659–667.

82 Kilpatrick LE, Lee JY, Haines KM, Campbell DE, Sullivan KE & Korchak HM (2002) A role for PKC-delta and PI 3-kinase in TNF-alpha-mediated antiapoptotic signaling in the human neutrophil. Am. J. Physiol. Cell Physiol. 283, C48–57.

83 Hauschild S, Tauber S, Lauber B, Thiel CS, Layer LE & Ullrich O (2014) T cell regulation in microgravity-\ The current knowledge from in vitro experiments conducted in space, parabolic flights and ground-based facilities. Acta Astronaut. 104, 365–377.

84 Li Q, Mei Q, Huyan T, Xie L, Che S, Yang H, Zhang M & Huang Q (2013) Effects of Simulated Microgravity on Primary Human NK Cells. Astrobiology 13, 703–714.

85 Paulsen K, Thiel C, Timm J, Schmidt PM, Huber K, Tauber S, Hemmersbach R, Seibt D, Kroll H, Grote K-H, Zipp F, Schneider-Stock R, Cogoli A, Hilliger A, Engelmann F & Ullrich O (2010) Microgravity-induced alterations in signal transduction in cells of the immune system. Acta Astronaut. 67, 1116–1125.

86 Zayzafoon M, Meyers VE & McDonald JM (2005) Microgravity: the immune response and bone. Immunol. Rev. 208, 267–280.

87 Gilis E, Gaublomme D, Staal J, Venken K, Dhaenens M, Lambrecht S, Coudenys J, Decruy T, Schryvers N, Driege Y, Dumas E, Demeyer A, De Muynck A, van Hengel J, Van Hoorebeke L, Deforce D, Beyaert R & Elewaut D (2019) MALT1-deletion in T-cells protects against the development of auto-immune arthritis, but causes spontaneous osteoporosis. Arthritis Rheumatol. Hoboken NJ.

88 Gao H, Liu H, Tang T, Huang X, Wang D, Li Y, Huang P & Peng Y (2018) Oleanonic acid ameliorates pressure overload-induced cardiac hypertrophy in rats: The role of PKCζ-NF-κB pathway. Mol. Cell. Endocrinol. 470, 259–268.

89 Islam SMA, Patel R & Acevedo-Duncan M (2018) Protein Kinase C-ζ stimulates colorectal cancer cell carcinogenesis via PKC-ζ/Rac1/Pak1/β-Catenin signaling cascade. Biochim. Biophys. Acta BBA – Mol. Cell Res. 1865, 650–664.

90 Staal J & Beyaert R (2018) Inflammation and NF-κB Signaling in Prostate Cancer: Mechanisms and Clinical Implications. Cells 7, 122.

91 Parker PJ, Justilien V, Riou P, Linch M & Fields AP (2014) Atypical protein kinase Cι as a human oncogene and therapeutic target. Biochem. Pharmacol. 88, 1–11.

92 Zhang S, Pan D, Jia X-M, Lin X & Zhao X (2017) The CARMA3-BCL10-MALT1 (CBM) complex contributes to DNA damage-induced NF-κB activation and cell survival. Protein Cell, 1–5.

93 Loh JT, Xu S, Huo JX, Kim SS-Y, Wang Y & Lam K-P (2019) Dok3-protein phosphatase 1 interaction attenuates Card9 signaling and neutrophil-dependent antifungal immunity. J. Clin. Invest. 130.

94 Eitelhuber AC, Warth S, Schimmack G, Düwel M, Hadian K, Demski K, Beisker W, Shinohara H, Kurosaki T, Heissmeyer V & Krappmann D (2011) Dephosphorylation of Carma1 by PP2A negatively regulates T-cell activation. EMBO J. 30, 594–605.

95 Okonechnikov K, Golosova O, Fursov M & Ugene team (2012) Unipro UGENE: a unified bioinformatics toolkit. Bioinforma. Oxf. Engl. 28, 1166–1167.

96 Edgar RC (2004) MUSCLE: multiple sequence alignment with high accuracy and high throughput. Nucleic Acids Res. 32, 1792–1797.

97 Guindon S, Delsuc F, Dufayard J-F & Gascuel O (2009) Estimating maximum likelihood phylogenies with PhyML. Methods Mol. Biol. Clifton NJ 537, 113–137.

98 Herrero J, Muffato M, Beal K, Fitzgerald S, Gordon L, Pignatelli M, Vilella AJ, Searle SMJ, Amode R, Brent S, Spooner W, Kulesha E, Yates A & Flicek P (2016) Ensembl comparative genomics resources. Database 2016, bav096.

99 Storz P, Döppler H & Toker A (2004) Protein kinase Cdelta selectively regulates protein kinase D-dependent activation of NF-kappaB in oxidative stress signaling. Mol. Cell. Biol. 24, 2614–2626.

100 Staal J, Alci K, De Schamphelaire W, Vanhoucke M & Beyaert R (2019) Engineering a minimal cloning vector from a pUC18 plasmid backbone with an extended multiple cloning site. BioTechniques 66, 254–259.

101 Wu EHT, Tam BHL & Wong YH (2006) Constitutively active alpha subunits of G(q/11) and G(12/13) families inhibit activation of the pro-survival Akt signaling cascade. FEBS J. 273, 2388–2398.

102 Schotte P, Denecker G, Van Den Broeke A, Vandenabeele P, Cornelis GR & Beyaert R (2004) Targeting Rac1 by the Yersinia effector protein YopE inhibits caspase-1-mediated maturation and release of interleukin-1beta. J. Biol. Chem. 279, 25134–25142.

103 Brinkman EK, Chen T, Amendola M & van Steensel B (2014) Easy quantitative assessment of genome editing by sequence trace decomposition. Nucleic Acids Res. 42, e168–e168.

104 Calcium phosphate-mediated transfection of eukaryotic cells (2005) Nat. Methods 2, 319–320.

105 Kimura A, Israël A, Le Bail O & Kourilsky P (1986) Detailed analysis of the mouse H-2Kb promoter: enhancer-like sequences and their role in the regulation of class I gene expression. Cell 44, 261–272.

106 Ishida T, Mizushima S, Azuma S, Kobayashi N, Tojo T, Suzuki K, Aizawa S, Watanabe T, Mosialos G, Kieff E, Yamamoto T & Inoue J (1996) Identification of TRAF6, a Novel Tumor Necrosis Factor Receptor-associated Factor Protein That Mediates Signaling from an Amino-terminal Domain of the CD40 Cytoplasmic Region. J. Biol. Chem. 271, 28745–28748.

107 Student (1908) The probable error of a mean. Biometrika 6, 1–25.

108 Wasserstein RL, Schirm AL & Lazar NA (2019) Moving to a World Beyond “p < 0.05.” Am. Stat. 73, 1–19.

109 Laemmli UK (1970) Cleavage of Structural Proteins during the Assembly of the Head of Bacteriophage T4. Nature 227, 680–685.

110 Bradford MM (1976) A rapid and sensitive method for the quantitation of microgram quantities of protein utilizing the principle of protein-dye binding. Anal. Biochem. 72, 248–254.

111 Tanghe G, Urwyler-Rösselet C, De Groote P, Dejardin E, De Bock P-J, Gevaert K, Vandenabeele P & Declercq W (2018) RIPK4 activity in keratinocytes is controlled by the SCFβ-TrCP ubiquitin ligase to maintain cortical actin organization. Cell. Mol. Life Sci. 75, 2827–2841.

112 Ran FA, Hsu PD, Wright J, Agarwala V, Scott DA & Zhang F (2013) Genome engineering using the CRISPR-Cas9 system. Nat. Protoc. 8, 2281–2308.

113 Baens M, Bonsignore L, Somers R, Vanderheydt C, Weeks SD, Gunnarsson J, Nilsson E, Roth RG, Thome M & Marynen P (2014) MALT1 auto-proteolysis is essential for NF-κB-dependent gene transcription in activated lymphocytes. PloS One 9, e103774.

